# From Decay to Rhythm: Coherent Biological Oscillators Require More Than Chemistry Alone

**DOI:** 10.1101/2025.04.15.648857

**Authors:** Yufei Wu, Sean X. Sun

## Abstract

Many biological systems exhibit sustained, coherent oscillations despite substantial noise. In contrast, chemical reaction systems governed by Markovian dynamics cannot sustain coherent ensemble oscillations, as the stochastic nature of state transitions inevitably causes the oscillation period to drift. To overcome this limitation, we propose a general mechanism that couples a Markovian system to at least one other degree of freedom, such as a mechanical system, to achieve noise-resistant coherent oscillations with a desired frequency. We introduce two approaches, targeting different dynamical modes in the Markovian system, and derive a governing principle for the non-Markovian system by analyzing the eigenvalues of the coupled dynamics. This principle is validated using a trimolecular reaction system, successfully producing sustained and coherent oscillations. Our study provides theoretical insights into how any chemical system can be coupled to another non-Markovian system to produce sustained and coherent oscillations with a precise period. We also make a fundamental observation that stability and control of stable limit cycles must arise from the non-Markovian part of the coupled system.

## Introduction

One of the hallmarks of biological systems is the presence of sustained, coherent, and robust periodic dynamics [1, 2, 3, 4]. Phenomena such as the heart beat [5], the cell cycle [6, 7], and the circadian rhythm [8, 9] are just a few examples where the periods of these collective oscillations are robust against external perturbations or changing environmental conditions. Recently, there have also been efforts to design oscillatory systems using tools of synthetic biology [10, 11]. Engineering these oscillatory systems and understanding natural biological oscillators require sound theoretical underpinning. Here, *coherence* is the persistence of oscillatory phase and period in time, as inferred from ensemble-averaged dynamics, typically quantified by a slow decay of autocorrelation functions or the narrowness of the power-spectral peak. Modeling of biological oscillators typically utilize kinetic rate equations describing multiple underlying chemical reactions and changes in densities of interacting components. For these types of models, conditions for establishing robust oscillatory dynamics are well known [12, 13]. However, it needs to be emphasized that kinetic rate equations (REs) are approximations to a more exact Markovian master equation description of kinetics [14]. The typical RE models that neglect fluctuations and higher order correlations are only first order moment approximations of Markovian dynamics. Markovian systems exhibit damped oscillations in the probability of occupying any given state. This oscillatory decay can be proven mathematically [15] and is an unavoidable consequence of the loss of phase coherence or oscillation amplitude, caused by the stochastic nature of state transitions [4, 16, 17, 18]. Previous studies have shown that the maximum phase coherence in such systems is bounded by the non-equilibrium driving force and the topology of the Markov network, as reflected in the finite number of coherent oscillations [19]. These results imply that robust biological oscillatory dynamics cannot arise from biochemical reactions alone. Other types of interactions are necessary to generate coherent oscillations or limit cycles in biological systems.

In this work, we explore the robustness of oscillations within the Markovian master equation formalism. For robust oscillations with a precise frequency, the ensemble average should exhibit sustained, undamped oscillations, whereas loss of coherence leads to damped decaying state probabilities. Previous studies have explored mechanisms for generating coherent oscillations, including positive and negative feedback, hysteresis [4], coupled oscillators [20, 21], and modifying driving force or network topology [19]. Here, we propose a novel mechanism to achieve sustained, coherent oscillations by coupling a Markovian system to additional unstable non-Markovian degrees of freedom (e.g., a mechanical system). This mechanism *does not* require coupling between oscillators. Instead, coupling to an external input is sufficient to generate coherent oscillation. We begin by demonstrating that coherent oscillations cannot emerge from Markovian dynamics. We then emphasize the distinction between the moment closure approach (i.e., REs) and the full Markovian dynamics as implemented by stochastic simulations. Using two auto-catalytic chemical reactions in both open and closed systems, we show that oscillations predicted by the moment closure approximation disappear when they are described more accurately by continuous-time Markovian dynamics. This discrepancy arises due to the truncation of higher-order moments in an expansion of the master equation. To achieve coherence, we propose two approaches targeting distinct dynamical modes in the Markovian system: direct decay and oscillatory decay. By analyzing the eigenvalues of the coupled Markovian and non-Markovian system, we derive a design principle for the non-Markovian component to enable sustained, coherent oscillations. This principle is validated using a trimolecular reaction system, where we successfully achieve coherent oscillations with the desired frequency. Our study enhances the understanding of coherent oscillations in biological systems, emphasizes the role of the non-Markovian component in oscillatory behavior, and provides theoretical insights into the design and control of stable and robust biological oscillators.

### Markov systems cannot develop coherent oscillations

For a continuous-time Markov chain, the time evolution of the probabilities of *n* distinct states is governed by the master equation:

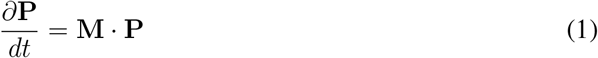

where **P**(*t*) is an *n*-dimensional vector denoting the probabilities of all *n* states at time *t*. The off-diagonal elements of the “transition matrix”, **M**_*βα*_ ≡ *k*_*α*→*β*_, is the probability per unit time to jump from the state *α* to *β*. The diagonal elements **M**_*αα*_ ≡ −*K*_*α*_ denotes the net escape rate from the state *α*. Because of the conservation of probability, all elements of **M** satisfy: ∑_*α*_ **M**_*αβ*_ = 0, and all diagonal elements are negative. Given this specific property, it can be proven that the real part of all non-zero eigenvalues of **M** are negative [15] (Supplemental Material, SM). This implies that **P** for any state must be a decaying function of time. Since the state probability represents ensembles of trajectories, each of which can individually oscillate, the reason for the decay is due to dephasing, or lack of phase coherence between trajectories. Thus, Markovian dynamics naturally cannot maintain an ensemble of coherent oscillators due to stochastic noise. This conclusion also holds regardless of whether the waiting time between jumps follows an exponential distribution characterized by a constant rate *k*_*α*→*β*_, or a more complex distribution.

#### Relationship between rate equations and the master equation

It has been noted that Markovian dynamics in Eq. 1 is the most fundamental description of chemical reactions [14]. A “state” in a reaction system is labeled by the number of various interacting species. Each reaction is a stochastic jump that changes the state or the number of reactant/products. This concept can be presented in a general form of the chemical master equation, described in [23]. We roughly sketch the theoretical framework here. Consider a chemical reaction network with *N* different molecule species and *R* chemical reactions. The reactions can be generally described by sets of chemical equations:

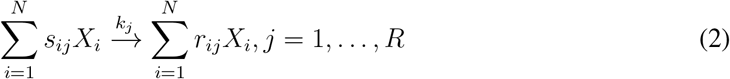

where *X*_*i*_ is the number of molecule *i, s*_*ij*_ and *r*_*ij*_ are the stoichiometric coefficients of reactants and products respectively. The chemical master equations can be written as:

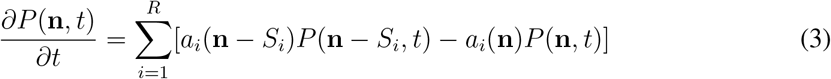

where *P* (**n**, *t*) is the probability of states with **n** molecules at time *t. a*_*i*_ is the propensity function of reaction *i*, which depends on the concentration of the reactants. *S* is the net stoichiometric matrix which is defined as: *S*_*ij*_ = *r*_*ij*_ − *s*_*ij*_, where *S*_*i*_ is the *i*^*th*^ column of matrix *S*. The chemical master equation is typically simulated using stochastic simulation algorithms (SSAs), such as the Gillespie algorithm [24, 25]. However, due to the high computational cost associated with large systems, an alternative method is the moment closure approximation [26], which analyzes statistical moments of the probability distribution rather than the full distribution itself (SM). This method truncates the hierarchy of moment equations at a finite order *q*. In particular, rate equations (REs) correspond to a first-order moment closure, capturing only the mean and neglecting fluctuations and higher-order correlations. REs can exhibit persistent oscillations. However, from the analysis above it is clear that the actual underlying Markov system cannot show coherent ensemble oscillations. Therefore, RE predictions of oscillations are misleading.

#### Archetypal chemical oscillators in open and closed systems

In this section, we present two examples that highlight the inability of Markov systems to generate coherent oscillations. We examine parameter regimes where oscillations are predicted by the approximate REs and compare RE solutions with those obtained from SSA.

A classic “chemical oscillator” is the Brusselator [27], a paradigmatic model of nonlinear autocatalytic reactions in open systems (Fig. 1a). This system involves two species, *X* and *Y*, and is governed by the following reactions:

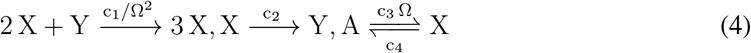

where *c*_1_, *c*_2_, *c*_3_, *c*_4_ are the reaction constants, and Ω is a system-size parameter. The concentration of molecule *A* is fixed, which is incorporated into the constant *c*_3_. Reaction 4 is solved using both the Gillespie algorithm and moment closure approximation (up to 5^*th*^ order). We compare the results of widely used REs (first-order moment closure) with stochastic simulation results for a small system size, where the initial molecule numbers are *X* = 1, *Y* = 1. In the parameter regime where oscillations are observed in REs, the ensemble-averaged dynamics obtained from stochastic simulations exhibit decay, as seen in the phase portraits of mean molecule numbers. Thus, RE approximation displays a limit cycle, whereas stochastic simulations converge to an attractor-like state (Fig. 1b). This decay is also evident in the time evolution of the ensemble-averaged molecule numbers (Fig. 1c). Higher-order moment closure schemes, such as the zero-closure approximation, yield more accurate results [26]. The origin of this decay lies in loss of coherence from stochasticity: although individual trajectories continue to oscillate with sustained amplitude and period, their phases gradually diffuse over time, as shown in single-trajectory dynamics (Fig. 1d). As a result, ensemble averaging leads to an apparent decay. The decay rate of the ensemble average therefore quantifies the loss of coherence, which is also reflected in the decaying autocorrelation function of the molecule number (Fig. 1e). An additional point of interest is whether moment closure approximations can accurately predict both the oscillation frequency and the decay rate with increasing order of closure. For the Brusselator, moment closure approximations of all orders provide relatively accurate predictions for the oscillation frequency but fail to predict the decay rate correctly (Fig. 1f). Notably, an accurate frequency prediction and converging but incorrect decay rate for REs are characteristics of open systems with an infinite supply of reactants. This behavior contrasts with what is observed in closed systems. The decaying property (loss of coherence) of oscillations is also examined in another example of a “biological oscillator” featuring the tumor suppressor P53, which involves feedbacks in a reaction network of an open system (SM, Fig. S2).

**Figure 1.**
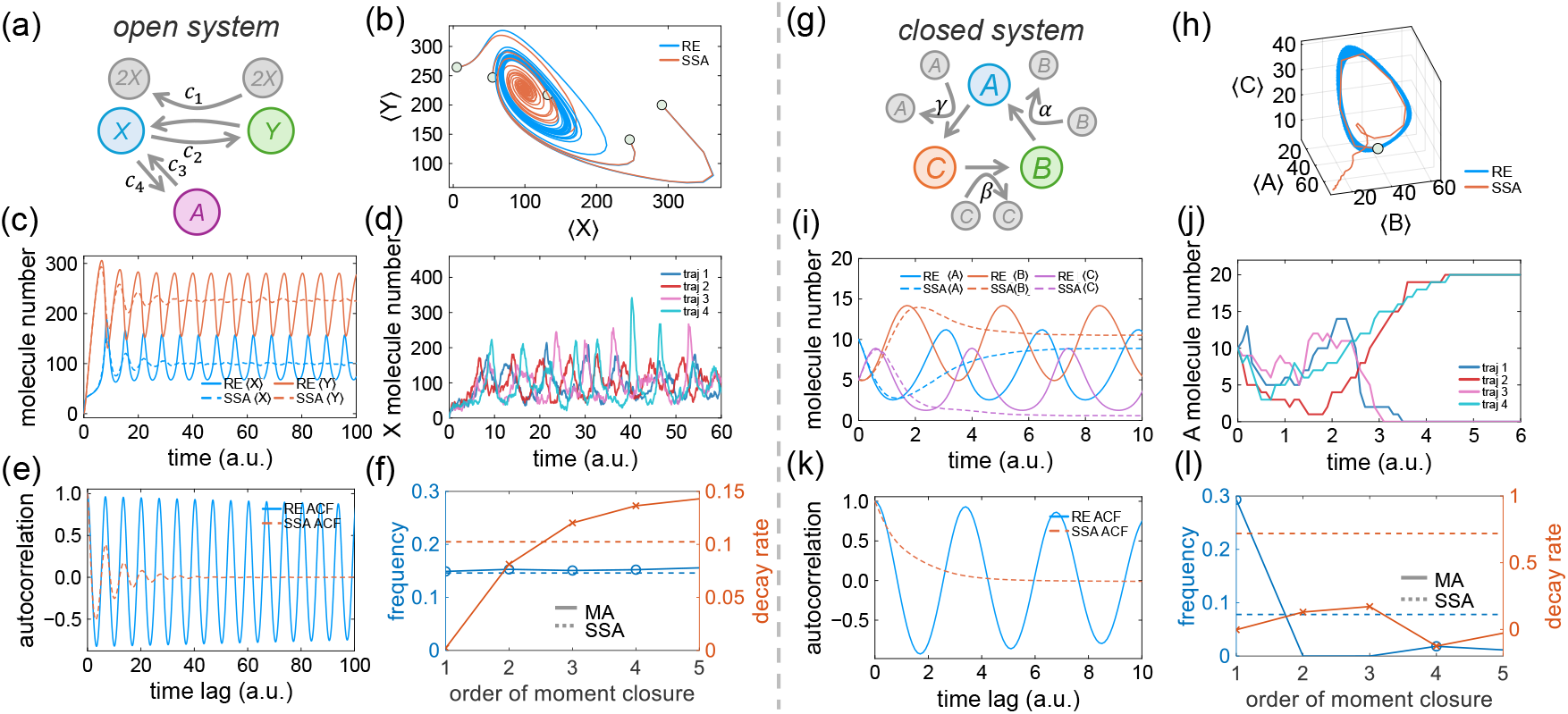
Stochasticity of Markovian systems prevents sustained coherent oscillations. (a) The Brusselator, an open-system autocatalytic chemical oscillator. (b) Phase portraits: rate equations (REs) exhibit a limit cycle, whereas stochastic simulation algorithm (SSA) results show decaying oscillations. (c) Ensembleaveraged time evolution of molecule numbers from REs and SSA. REs (first-order moment closure) predict sustained oscillations in the ensemble average but fail to capture the decay observed in SSA, indicating loss of coherence. (d) Representative individual SSA trajectories of molecule-number dynamics of X. (e) Autocorrelation function computed from the molecule number of X. (f) Frequency and decay rates derived from various orders of moment closure approximations (MAs) and SSA simulations. MAs accurately predict oscillation frequencies but fail to capture decay rates. (g) Trimolecular autocatalytic reaction in a closed system. (h) Phase portraits: SSA shows convergence to a single-species steady state. (i) Ensemble-averaged molecule-number dynamics from RE and SSA; decay in this closed system arises from loss of sustained oscillations in individual trajectories. (j) Representative individual SSA trajectories of molecule-number dynamics of A. (k) Autocorrelation function computed from the molecule number of A. (l) Frequency and decay rates derived from MAs and SSA simulations, highlighting the inability of MAs to accurately approximate either property in a closed system. The “central-moment-neglect” scheme is used for moment closure [22]. In panels b, c, h, and i, each SSA trajectory represents the time evolution averaged over 1000 independent realizations with the same initial condition.

The previous example explores an open system with infinite pool of reactants. However, biological systems are finite and cells have a limited number of interacting proteins. To investigate this, we examine a minimal closed system exhibiting oscillatory behavior described by REs. This system involves three species (Fig. 1g), with the following reactions:

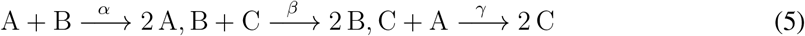

where *k*_1_, *k*_2_, and *k*_3_ represent the reaction rate constants. In this closed system, the total number of *A, B*, and *C* remains constant over time. We simulate this system starting with the state *n*_*A*_ = 10, *n*_*B*_ = 5, *n*_*C*_ = 5. Fig. 1h-i shows that again, although REs predict sustained oscillation, the actual dynamics exhibit decay in the ensemble-averaged molecule number. However, this behavior differs from that of the Brusselator, an open system. In the trimolecular closed system considered here, individual trajectories do not exhibit oscillations with constant amplitude (Fig. 1j). The observed decay is a characteristic feature of small, closed systems with fixed total molecule numbers and only forward reactions. In such systems, the dynamics can readily reach absorbing states in which only one molecular species remains. Consequently, the steady-state probability distribution becomes concentrated on a small number of discrete states, each corresponding to a single dominant species. This characteristic renders the commonly used moment closure schemes ineffective, as the probability distribution is not accurately captured by analytical distribution functions. Consistent with this picture, the autocorrelation function also exhibits decay (Fig. 1k). Unlike the Brusselator, rate equation approximations for this closed system fail to accurately predict either the oscillation frequency or the decay rate (Fig. 1l).

The observations in closed systems are particularly relevant for synthetic biology, where genetic and biochemical circuits operate in confined cell volumes with finite numbers of molecules and substantial intrinsic noise. Synthetic oscillators, such as transcriptional clocks and repressilator type networks, therefore more closely resemble the finite, stochastic systems studied here than idealized infinite-reservoir models [28, 29]. In this regime, RE descriptions can predict oscillatory behavior but fail to capture phase diffusion, loss of temporal coherence, and decay observed in stochastic dynamics. These examples demonstrate that purely Markovian chemical systems cannot generate coherent oscillations at the ensemble-averaged level. Apparent sustained oscillations in simplified Markov models arise from moment-closure approximations that neglect higher-order moments. This raises two closely related questions: how do biological systems, such as circadian clocks, achieve sustained and temporally coherent oscillations with precise and robust frequencies despite intrinsic noise, and, in synthetic biology, what engineering principles can be employed to design circuits that generate similarly robust and coherent oscillations?

### Coupling a Markov system with an unstable non-Markov system generates coherent oscillations

In order to achieve coherent oscillation, it is necessary to introduce another degree of freedom that influences the probability of each state to mitigate the decay in Markov systems. In this study, we show a specific realization by coupling the stable Markov system to an unstable non-Markov system (Fig. 2a) (a control input). This type of models appears in many multi-physics models of biological systems such as the Hodgkin-Huxley model [30], which couples ion channel opening probabilities with cell membrane voltage. While ion transport via ion channels follows a Markov process described by the master equation, the electrical potential is governed by membrane capacitance and therefore does not follow Markovian dynamics (SM).

**Figure 2.**
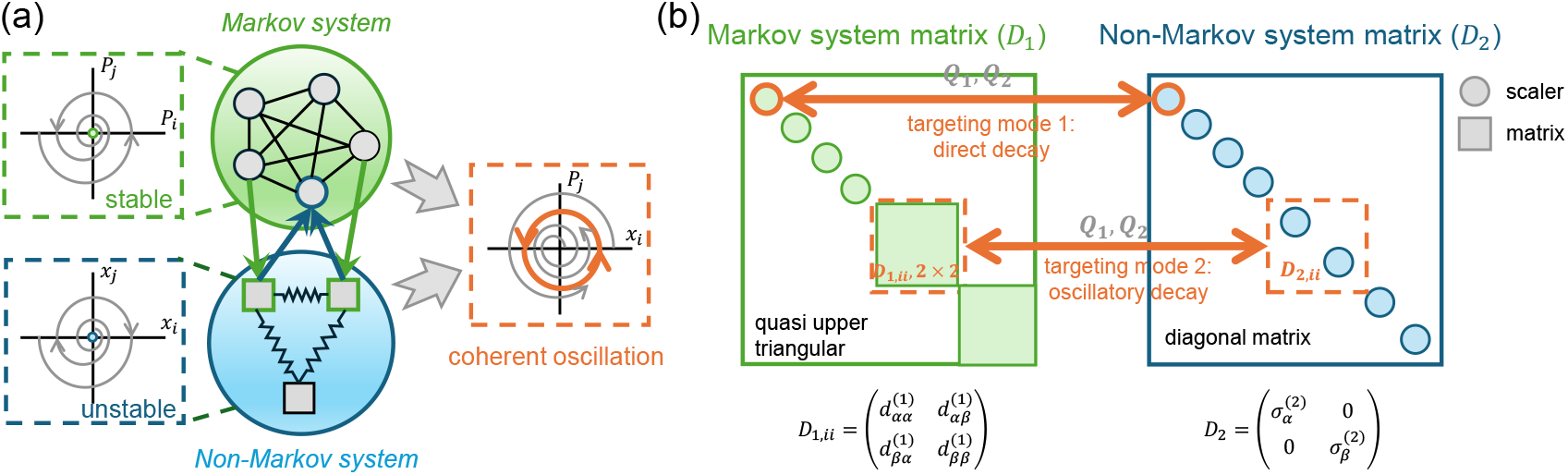
(a) Coupling a stable Markov system to an unstable non-Markovian system can generate coherent ensemble oscillations. (b) Schematic representation of the coupling design between a non-Markov system and a Markov system targeting different dynamical modes. When the diagonal block of the non-Markov system corresponds to a scalar element in the Markov matrix **D**_1_, the targeted mode represents direct decay, and coupling transforms this mode into sustained oscillations. Similarly, when the diagonal block of the non-Markov system corresponds to a 2 *×* 2 submatrix in **D**_1_, the coupling converts a decaying oscillatory mode into coherent oscillations. Coupling between the two systems is facilitated through the coupling coefficients 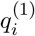 and 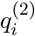.

In the most general form, we can write the linearized governing equation for the coupled system as (SM):

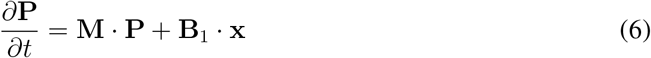

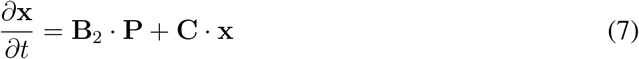

where **P** is a *n* dimensional vector that denotes the probability of each state in the Markov system while **x** is a *l* dimensional vector which is the state variable of the coupled system (e.g., a mechanical system). Consequently, the coupling matrices **B**_1_ and **B**_2_ are with dimensions *n × l* and *l×n*. In order to maintain probability conservation, **B**_1_ should satisfy: ∑_*i*_ **B**_1,*ij*_ = 0. The matrices (**M, B**_1_, **B**_2_, **C**) forms an overall (*n* + *l*) *×* (*n* + *l*) matrix **A**. An analytical solution for the input variable **x** can be expressed in terms of **P** as:

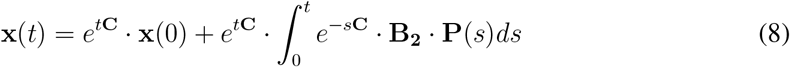

We see that **x**(*t*) is an integral of the probability **P**(*t*), which is also the concept behind an integral controller [31]. For the purpose of generating coherent ensemble oscillations, we aim to design the system so that the state probabilities exhibit sustained oscillations without decay. We seek to design coupling matrices **B**_1_, **B**_2_, such that (**A**) has at least one eigenvalue with zero real part and non-zero imaginary part. To prevent instability, we also require that the real parts of other eigenvalues are non-positive. Denoting the new eigenvalues of **A** as *λ*, the characteristic equations are:

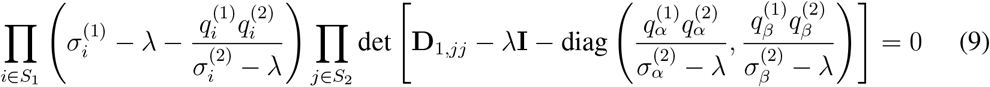

Here 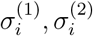 are the real eigenvalues of matrices **M** and **C**. 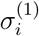 excludes the inherent zero eigenvalue in **M. D**_1_ is a quasi-upper triangular matrix of **M** after Schur decomposition. The diagonal blocks are either scalar or 2 by 2 matrices. For convenience, all matrix blocks are placed at the bottom right, denoted by 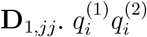 describes the coupling strength between the Markov and non-Markov systems. Detailed derivation is in the SM. *S*_1_ is the set of positional indices of scalar diagonal elements in **D**_1_, *S*_2_ is the set of positional indices corresponding to 2*×*2 blocks in **D**_1_. *α, β* are the indices of individual scalar entries corresponding to the *j*^*th*^ block.

From Eq. 9, we observe two approaches that can both achieve pure imaginary eigenvalues corresponding to coherent oscillations. The first approach targets the real eigenvalues of the Markov system (first product in Eq. 9), while the second targets the complex eigenvalues (second product, Fig. 2b). These approaches lead to different dynamical transitions. The first approach turns a stable non-oscillatory mode into a sustained oscillatory mode (coherent oscillation), while the second turns a decaying oscillatory mode into a sustained one. For the first approach, the condition for eigenvalues with zero real part is 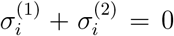 and 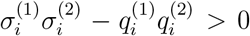. For the other eigenvalues, the real part of *λ* should be non-positive. The frequency of the coupled system is 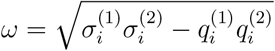.

The condition for coherent oscillation can be interpreted as follows: Since the Markovian system always has negative real eigenvalues 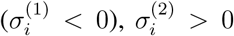 must hold, implying that the non-Markovian system should be inherently unstable. Additionally, 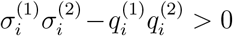 requires 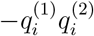 o be large enough to make this term positive. This means that: 1. There must be strong coupling between the two systems. 2. The systems must influence each other in opposite directions (e.g., A inhibits B while B enhances A), i.e., a negative feedback loop, a common feature observed in oscillatory systems [32, 33]. When the non-Markovian system targets the complex eigenvalues of the Markov matrix, the second product in Eq. 9 must generate pure imaginary roots. Denoting individual elements of **D**_1,*ii*_ as *d*_*αβ*_ (*i* ∈ *S*_2_), the new eigenvalues *λ* of the coupled system can be solved from:

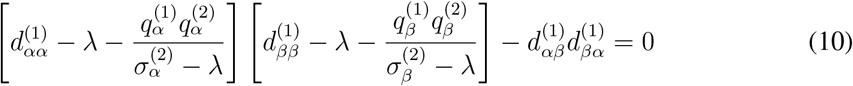

Eq. (10) is a quartic equation with four roots. Assuming the *β*^*th*^ element in the non-Markovian system is weakly coupled to the Markov system, we can approximate 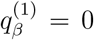. This decouples one dimension, reducing the quartic equation to a cubic of the form *aλ*^3^ + *bλ*^2^ + *cλ* + *d* = 0. For the equation to have pure imaginary roots, the necessary and sufficient conditions are *bc* = *ad* and *b >* 0, *c >* 0.

#### Example - generating coherent oscillation in a tri-molecular reaction

We use a trimolecular reaction as an example to design a coupled system that generates sustained and coherent ensemble oscillations (Fig. 3a, Fig. S3). The coupled system is described by Eqs. 6-7. For illustration, we design a simple one-dimensional input, *x*(*t*), targeting the largest nonzero real eigenvalue of matrix **M**. The dynamical results are shown in Fig. 3b. Coupling with a non-Markovian system can generate coherent oscillations and transition from a decaying to an oscillatory dynamical mode, with sustained phase coherence evidenced by a persistent autocorrelation function (Fig. 3c).

**Figure 3.**
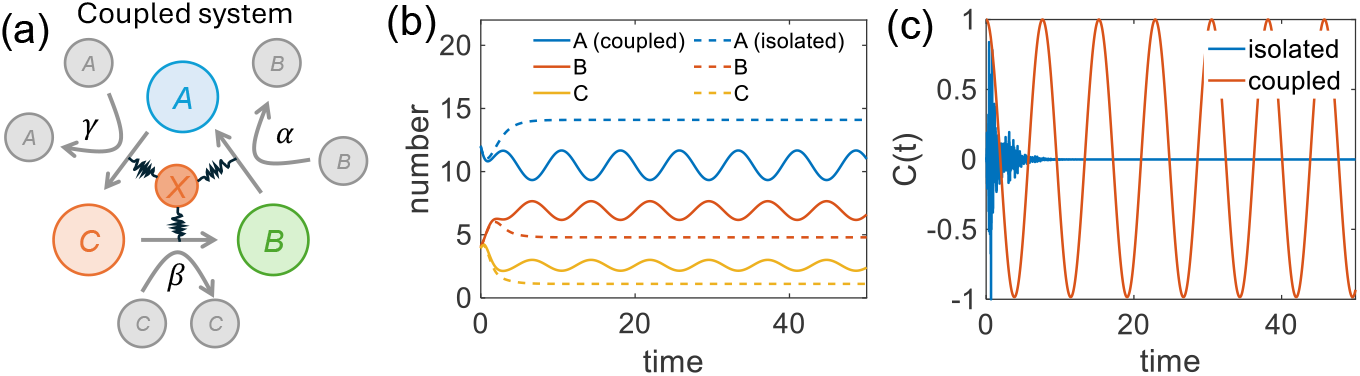
Generation of coherent oscillations in a trimolecular reaction system. (a) Schematic representation of the coupled oscillatory system. (b) Time evolution of molecular populations in the isolated Markov system vs. the coupled system. (c) Autocorrelation functions in the isolated Markov system and the coupled system. The autocorrelation function is normalized to range between −1 and 1.

Results show that it is possible to design an oscillator with a targeted frequency by adjusting the eigenvalue of the coupled system 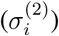 and coupling coefficients 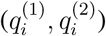 according to Eq. As discussed previously, the Markovian system has two dynamical modes: a direct decaying mode (real eigenvalue) and an oscillatory decaying mode (complex eigenvalues). Targeting the direct decaying mode in the non-Markovian system transforms the decaying process into a coherent oscillation. The obtained frequency as a function of 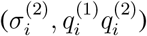 is illustrated in a phase diagram (Fig. 4a) with four regions (I-IV). These regions represent distinct dynamical phases of the coupled system, and at the phase boundaries correspond to phase-transitions where the system switches abruptly between qualitatively different dynamics (stable decay, coherent oscillation, unstable oscillation, etc.). Generating coherent ensemble oscillations requires precise eigenvalue design, as the system becomes either stable or unstable when the eigenvalue is too small or too large. The coupling effects 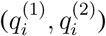 further categorize the system into oscillatory or non-oscillatory states. A strong negative coupling between the systems is required for coherent oscillations, as outlined in the previous section. This requirement is also reflected in the oscillation frequency near the coherent regime as a function of coupling strength, which exhibits a phase transition with square-root (1/2) scaling (Fig. 4b). Fig. S4 demonstrates the accuracy of our design within a target frequency.

**Figure 4.**
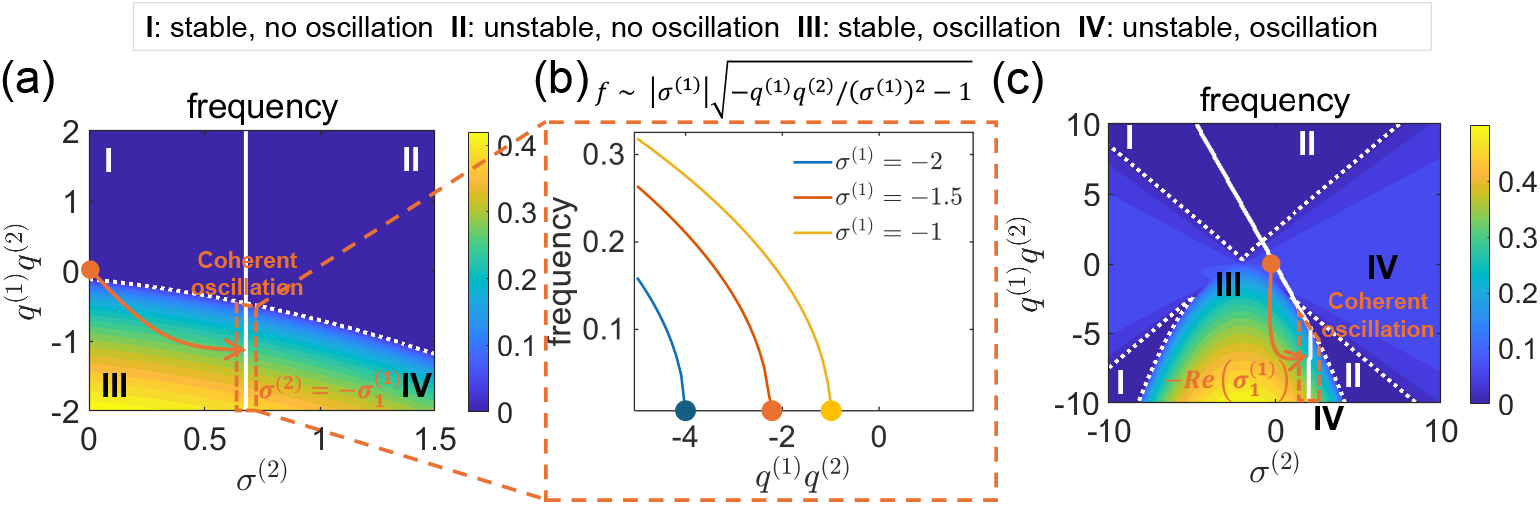
(a) Phase diagram showing system frequency as a function of the non-Markov eigenvalue (*σ*^(2)^) and coupling (*q*^(1)^*q*^(2)^), targeting a directly decaying Markov mode. Arrow indicates transition to coherent oscillations. (b) Phase transition in oscillation frequency versus coupling strength for a directly decaying Markov mode, exhibiting 1/2 scaling behavior. (c) Phase diagram targeting an oscillatory Markov mode; arrow shows transition from decaying oscillations to sustained oscillations.

Similarly, we can obtain a frequency phase diagram when the non-Markov system targets the oscillatory decaying mode in the Markov system (corresponding to complex eigenvalues). This transforms a decaying oscillatory mode to a sustained coherent oscillatory mode. Here, we take the assumption made in the previous section that the two by two matrix in the non-Markov system corresponding to the targeting mode in Markov system has one eigenvalue much larger than the other, denoted by *σ*^(2)^. A more complex phase diagram is seen in Fig. 4c. A square-root (1/2) scaling behavior, similar to that observed when targeting the directly decaying mode, is also found near the coherent oscillation regime (Fig. S5).

When targeting the oscillatory decay mode, in addition to designing a system with a preferred frequency, we can also eliminate certain frequency components from the original Markov system (Fig. 4c). For example, we can introduce a positive coupling (*q*^(1)^*q*^(2)^ *>* 0) to change an oscillating mode in the original Markov system to a non-oscillatory one (Upper Regions I&II).

#### Generating biological limit cycles

It is important to note that strictly speaking, coherent oscillations generated by Eqs. (6-7) are not true limit cycles, i.e., stable trajectories in phase space that attract nearby trajectories. A necessary condition for a limit cycle is nonlinearity in Eqs. (6–7). Since Markovian chemical systems are intrinsically linear, a possible design principle is for the non-Markovian system to provide the source of stability — for instance, by introducing a cubic term into the equation for **x** in Eq. (7). This is also what is observed in the Hodgkin-Huxley model.

## Conclusion and discussion

In this paper, we demonstrate that the Markovian master equation, commonly used to model chemical reaction systems, cannot generate coherent ensemble oscillations. Here, “coherent oscillations” refers to both sustained amplitude and temporal coherence among trajectories. While many studies acknowledge the decaying nature of Markovian dynamics, they often neglect it, which is reasonable for short time scales. However, in processes such as circadian rhythms and cell cycle progression, where the time scale is much longer, we show that an additional non-Markovian input is necessary to prevent the loss of phase coherence, ensuring a precise oscillation period. One such example is the Hodgkin-Huxley model, which couples ion channel dynamics (Markovian) to membrane voltage dynamics (non-Markovian), enabling the generation of sustained and coherent voltage oscillations [34]. Analogous to electrical excitability in neurons, sustained and coherent oscillations are also observed in the cell cortex. These oscillations arise from coupling between Markovian chemical processes, such as actin polymerization and myosin recruitment, and non-Markovian mechanical variables including cortical and membrane tension. This mechanochemical feedback allows actomyosin activity to generate mechanical forces, while mechanical stress in turn regulates actomyosin dynamics, producing sustained coherent oscillations in contractile forces and propagating waves in tension and signaling molecules such as Ca^2+^ [35, 36, 37]. At larger scales, similar mechanochemical couplings in epithelial tissues lead to oscillations in cell volume and shape [38, 39]. Similarly, in synthetic biology, designing biological circuits capable of sustaining coherent oscillations remains an active area of research [11, 40] and our work shows that additional design criterion is desirable. Our work identifies the strategy for a chemical system coupled to another degree of freedom (Eq. 6-7). Designing such coupled systems may benefit from optimal control strategies that minimize the error between the predicted and targeted oscillatory trajectories with minimum dissipation [41], ensuring precise control of oscillation period and amplitude. Advanced AI-driven tools, such as neural ODEs in optimal control [42], provide powerful methods for handling time-varying target signals in oscillator design, especially for systems with unknown eigenvalues and eigenvectors.

Our work focuses on the temporal coherence of an ensemble of oscillators and their ability to maintain a well-defined and precise period despite the presence of noise. In contrast, a substantial body of research concerns synchronization in populations of oscillators, where the objective is not only for each oscillator to maintain a coherent period but also for the entire group to align their phases through coupling or shared signals. Synchronization is therefore fundamentally different from temporal coherence: coherence pertains to the stability of an individual oscillator’s period, whereas synchronization requires both period matching and phase alignment among many oscillators. It is possible, for example, for multiple oscillators to exhibit strong temporal coherence—each maintaining a precise period—while starting with different phases and preserving those phase differences over time; such a system would be coherent but not synchronized. A typical example is the circadian oscillator in cyanobacteria, which exhibits robust oscillations in individual cells while lacking phase synchronization even among neighboring cells [43]. Because synchronization imposes constraints on the relative phases of oscillators in addition to their periods, it is generally considered a stronger condition than individual-oscillator coherence. Our study contributes to understanding the mechanisms that maintain temporal coherence at the single-oscillator level, forming a foundation for future exploration of synchronization in coupled oscillator networks governed by Markovian dynamics.

Ultimately, biological oscillators are limit cycles, as they not only maintain a desired frequency but also enforce a fixed amplitude, which is essential for biological functions like heartbeat regulation. Additionally, robustness of biochemical systems to intrinsic and extrinsic noise is crucial for maintaining an accurate period, which depends on the structure of the chemical reaction network and the transition rates at each step [44, 45]. while our model accounts for intrinsic noise (copy number fluctuations) through the chemical master equation, we do not consider extrinsic noise [46]. For systems exhibiting limit cycles, small perturbations due to extrinsic noise should dissipate over time, ensuring robust oscillatory behavior. Future work should incorporate both nonlinearity and extrinsic noise to study the robustness of oscillations under real-world perturbations.

## Acknowledgments

This work has been funded in part by NIH R01GM134542. We would like to thank Noah J. Cowan for helpful discussions.

