## Supplementary Note for "From Decay to Rhythm: Coherent Biological Oscillators Require More Than Chemistry Alone"

#### Supplementary note 1 - A mathematical proof that continuous-time Markov chain cannot generate coherent oscillation

The proof uses Perron-Frobenius theorem [1], which states: For a non-negative matrix $\mathbf{A}$, its spectral radius $\rho(\mathbf{A})$ is an eigenvalue of $\mathbf{A}$ and there is a non-negative non-zero vector $\mathbf{x}$ such that $\mathbf{Ax} = \rho(\mathbf{A})\mathbf{x}$. In the master equation, we define a new matrix as follows: Let $a = \max(\text{abs}(\mathbf{M}_{\alpha\alpha}))+\delta$, where $\delta > 0$. Let $\mathbf{A} = \mathbf{M}+a\mathbf{I}$, where $\mathbf{I}$ is the identity matrix. The newly defined matrix $\mathbf{A}$ is non-negative and according to Perron-Frobenius theorem, the spectral radius $\rho(\mathbf{A})$ is one of the eigenvalues of matrix $\mathbf{A}$ and all other eigenvalues are less or equal than $\rho(\mathbf{A})$ in magnitude. We denote the eigenvalues of $\mathbf{A}$ as $\lambda$ and eigenvalues of $\mathbf{M}$ as $r$. The maximum eigenvalue is denoted as $\lambda_m$ for matrix $\mathbf{A}$ and the corresponding eigenvector is $\mathbf{x_m}$, which is non-zero and non-negative. According to the definition of matrix $\mathbf{A}$, $r_m = \lambda_m - a$ is an eigenvalue of matrix $\mathbf{M}$ and it shares the same eigenvector $\mathbf{x}_m$. The algebraic multiplicity of $\lambda_m$ can be larger than one, i.e., there could be multiple eigenvalues with the same value $\lambda_m$. However, all of them will be real. We denote all other eigenvalues of matrix $\mathbf{A}$ as $\lambda_r$, we will have $\lambda_r < \lambda_m$ for real numbers. For complex eigenvalues, we have $\text{real}(\lambda_r) < |\lambda_r| <= \lambda_m$.

Matrix $\mathbf{M}$ satisfies the equation: $(\mathbf{M}+a\mathbf{I})\mathbf{x}_m = \lambda_m\mathbf{x}_m$, which can be rewritten as: $\mathbf{M}\mathbf{x}_m = (\lambda_m - a)\mathbf{x}_m$. We then sum over all the elements in the vector: $\sum_\alpha \sum_\beta M_{\alpha\beta} x_{m,\beta} = (\lambda_m - a)\sum_\beta x_{m,\beta}$. Given the specific property of transition matrix, $\sum_\alpha M_{\alpha\beta} = 0$, the left hand side is zero. On the right hand side, since $\mathbf{x_m}$ is non-negative and non-zero, $\sum_\beta x_{m,\beta} > 0$. Therefore, $r_m = \lambda_m - a = 0$. Note that is also the maximum eigenvalue of matrix $\mathbf{M}$. Since we care about the oscillation regime, we only care about complex eigenvalues. All complex eigenvalues of $\mathbf{M}$ can be expressed as $r = \lambda - a$. Therefore, $\text{real}(r) = \text{real}(\lambda) - a < \lambda_m - a = 0$. This proves that for all complex eigenvalues of transition matrix $\mathbf{M}$, the real parts are negative, so sustained oscillations in probability dynamics cannot arise from standard continuous-time Markov chains.

Note that the decay of the probability dynamics does not necessarily imply that individual oscillators stop oscillating. One possibility is that individual trajectories cease to exhibit sustained oscillations. Alternatively, individual trajectories may maintain their oscillation amplitude, but their periods are continually perturbed by stochastic fluctuations, leading to a gradual loss of phase coherence. As a result of ensemble averaging, this loss of coherence manifests as an apparent decay of oscillation amplitude. The same effect is captured by the autocorrelation function, $C(\tau) = \langle n(t)n(t+\tau)\rangle$, where $n(t)$ denotes the number of a given molecular species at time $t$, and $C(\tau)$ similarly decays over time. In the examples presented in the main text, the Brusselator exhibits sustained oscillations in individual trajectories but a loss of coherence over time. In contrast, for the trimolecular autocatalytic reaction, individual trajectories cannot sustain oscillations due to the closed-system nature and low molecule-number constraints (Fig. 1). In both cases, autocorrelation functions obtained from stochastic simulations (SSA) exhibit decay, arising either from loss of phase coherence or from the inability to sustain oscillations with constant amplitude. By contrast, rate equation approximations fail to capture either the loss of coherence or the decay of oscillation amplitude (Fig. S1).

### Supplementary note 2 - Relationship between moment closure approximation and the master equation

It has been noted that Markovian dynamics in Eq. 1 is the most fundamental description of chemical reactions [2]. A "state" in a reaction system is labeled by the number of various interacting species. Each reaction is a stochastic jump that changes the state or the number of reactant/products. This concept can be presented in a general form of the chemical master equation, described in [3]. We roughly sketch the theoretical framework

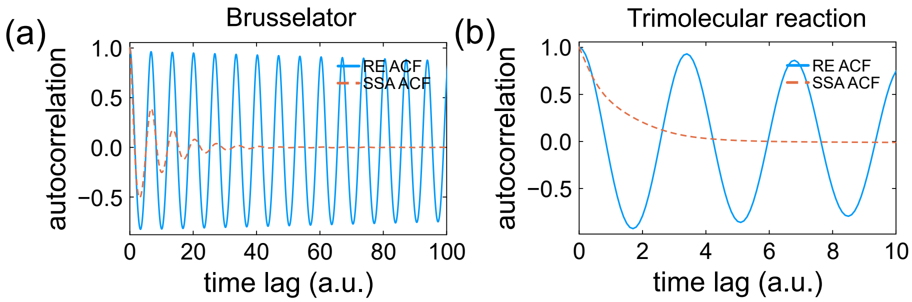


Fig. S1: Autocorrelation functions of (a) the Brusselator (calculated from molecule number of $X$) and (b) the trimolecular autocatalytic reaction (calculated from molecule number of $A$). The autocorrelation functions are normalized to range between −1 and 1.

here. Consider a chemical reaction network with $N$ different molecule species and $R$ chemical reactions. The reactions can be generally described by sets of chemical equations:

$$\sum_{i=1}^{N} s_{ij} X_i \xrightarrow{k_j} \sum_{i=1}^{N} r_{ij} X_i, j = 1, \ldots, R \tag{1}$$

where $X_i$ is the number of molecule $i$, $s_{ij}$ and $r_{ij}$ are the stoichiometric coefficients of reactants and products respectively. The specific form of chemical master equations can be written as:

$$\frac{\partial P(\mathbf{n}, t)}{\partial t} = \sum_{i=1}^{R} [a_i(\mathbf{n} - S_i) P(\mathbf{n} - S_i, t) - a_i(\mathbf{n}) P(\mathbf{n}, t)] \tag{2}$$

where $P(\mathbf{n}, t)$ is the probability of finding molecules with numbers $\mathbf{n}$ at time $t$. $a_i$ is the propensity function of reaction $i$, which depends on the concentration of the reactants. $S$ is the net stoichiometric matrix which is defined as: $S_{ij} = r_{ij} - s_{ij}$, where $S_i$ is the $i^{th}$ column of matrix $S$. The standard approach for simulating the chemical master equation is the stochastic simulation algorithm (SSA), such as the Gillespie algorithm [4]. However, due to the high computational cost associated with large systems, an alternative method is the moment closure approximation [5]. In this approach, instead of considering the full probability of each state, only moments of the probability distribution are analyzed. The moment equations can be written as:

$$\frac{d\mu_i}{dt} = \sum_{\boldsymbol{n}} n_1^{i_1} \ldots n_N^{i_N} \frac{dP(\mathbf{n}, t)}{dt} = \sum_{r} \sum_{|\boldsymbol{j}|=0}^{|\boldsymbol{i}|-1} \prod_{k=1}^{N} \binom{i_k}{j_k} S_{kr}^{i_k - j_k} \langle a_r(\boldsymbol{n}) \prod_{l=1}^{N} n_l^{j_l} \rangle \tag{3}$$

Here, $\mu_i$ represents the $i^{\text{th}}$-order moment: $\mu_i = \langle \prod_{k=1}^{N} n_k^{i_k} \rangle$. The multi-index notation is expressed as: $\sum_{|\boldsymbol{j}|=0}^{|\boldsymbol{i}|-1} = \sum_{0 \le j_l \le i_l (l=1,\ldots,N), 0 \le \sum_{l=1}^{N} j_l \le |\boldsymbol{i}|-1}$. Details of the derivation and implementation are provided in reference [6, 7].

The moment equations are nonlinear ODEs. To close the series of moment equations, it must be truncated at a finite order $q$. The choice of closure depends on the system, common options including zero closure, normal closure, log-normal closure, among others [5]. Notably, the typical REs correspond to the moment closure scheme truncated at the first order, or the mean of the full probability distribution. This implies that REs generally neglect fluctuations and higher order correlations.

#### Supplementary note 3 - P53 network

The decay in probability dynamics is further illustrated using another example of a "biological oscillator" in an open system. This example focuses on the molecule $P53$, whose oscillation frequency resembles that of the cell cycle. Three molecules are involved: p53, Mdm2 precursor, and Mdm2. Unlike the Brusselator system discussed earlier, the p53-Mdm2 system involves non-polynomial propensities and incorporates the effect of negative feedback. Details of the reaction network are provided in [8]. A schematic illustration of the system is shown in Fig. S2(a). The number of these molecules (p53, Mdm2 precursor, and Mdm2) are denoted by $X, Y_0, Y$, respectively. This is an open system, where P53 is communicating with an infinite source. Specifically, there is a negative feedback of Mdm2 ($Y$) on the net production of P53 ($X$) by enhancing its degradation. The reactions can be written in several chemical equations:

$$\mathrm{A} \underset{\mathrm{k_2}}{\overset{\mathrm{k_1}}{\rightleftharpoons}} \mathrm{X}, \mathrm{X} \xrightarrow{\mathrm{k_3XY/(X+k_7)}} \mathrm{A}, \mathrm{X} \xrightarrow{\mathrm{k_4}} \mathrm{Y_0}, \mathrm{Y_0} \xrightarrow{\mathrm{k_5}} \mathrm{Y}, \mathrm{Y} \xrightarrow{\mathrm{k_6}} \mathrm{B} \tag{4}$$

where $k_1 - k_7$ are the reaction constants. Molecule $A$ and $B$ has fixed concentration. Reaction 4 is solved in both approaches of Gillespie algorithm and moment closure approximation (up to $5^{th}$ order). As shown in Fig. S2(b), this biological oscillator exhibits decay in the ensemble-averaged dynamics, indicating a loss of coherence and period precision. This behavior is clearly reflected in individual trajectories obtained from stochastic simulations (Fig. S2(c)) and in the decay of the autocorrelation function calculated from the molecule number of P53 (Fig. S2(e)). However, sustained and coherent oscillations are still observed in the rate equation approximation (Fig. S2(b)(d)). This decaying oscillation is in contrast with the long lasting coherent oscillation in real biological systems. This discrepancy suggests that real biological systems are not purely Markovian systems governed by the typical master equation. Instead, additional mechanisms or couplings beyond standard Markovian dynamics are likely required to sustain coherent oscillatory behavior in these systems. Similar to the Brusselator example, we can approximate the frequency by moment closure approximation (with normal closure scheme) and we can obtain similar results as the Brusselator (Fig. S2(f)).

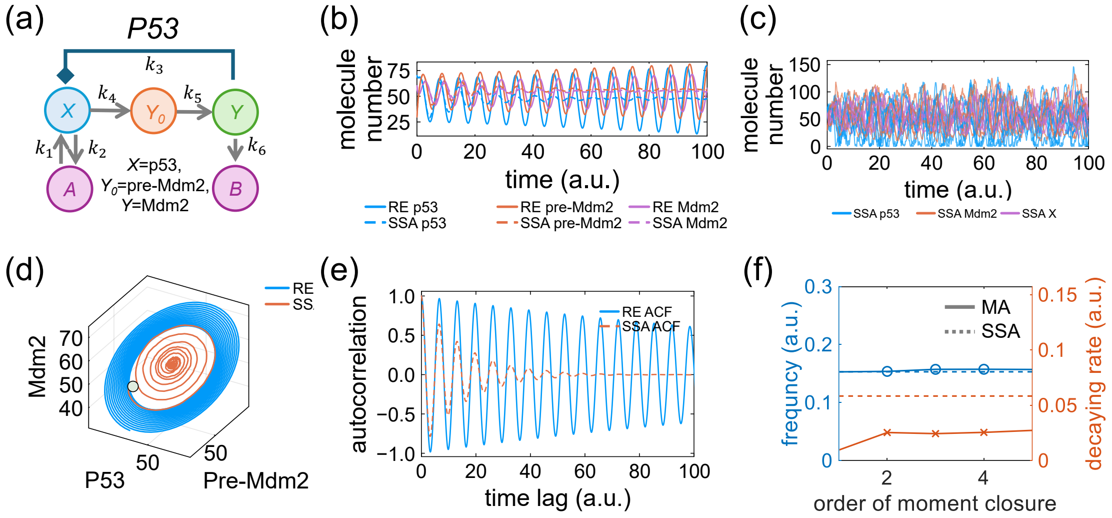


Fig. S2: p53 Network: A canonical biological oscillator governed by negative feedback. (a) Schematic representation of the reaction. Variables $X, Y_0, Y$ correspond to the number of p53, Mdm2 precursor, and Mdm2, respectively. (b) Temporal dynamics of molecular populations. (c) Representative individual SSA trajectories. (d) Phase portraits depicting the molecular dynamics. (e) Autocorrelation function calculated from the molecule number of p53. The autocorrelation function is normalized to range between -1 and 1. (f) Frequency and decay rate estimations obtained from moment closure approximations of varying orders and stochastic simulation algorithm (SSA) results.

#### Supplementary note 4 — Moment dynamics of the trimolecular reaction network

In this part, we derive the moment equations of the trimolecular reaction in the main text and show that typical chemical oscillations are a byproduct of truncation error in calculating moments in the master equation. In this trimolecular system, there are three coupled reactions:

$$A + B \to 2A(\alpha) \tag{5}$$

$$B + C \to 2B(\beta) \tag{6}$$

$$C + A \to 2C(\gamma) \tag{7}$$

We first consider the kinetic equations based on the law of action of mass. We can write down three ODEs describing the dynamics of concentrations of the three molecules $A, B, C$:

$$\frac{d[A]}{dt} = \alpha[A][B] - \gamma[A][C] \tag{8}$$

$$\frac{d[B]}{dt} = \beta[B][C] - \alpha[A][B] \tag{9}$$

$$\frac{d[A]}{dt} = \gamma[A][C] - \beta[B][C] \tag{10}$$

where $\alpha, \beta, \gamma$ are the rate coefficients of the three reactions respectively. These ODEs can be numerically solved and generates sustained coherent oscillation, and the results are shown in Fig. 1(f) in the main text.

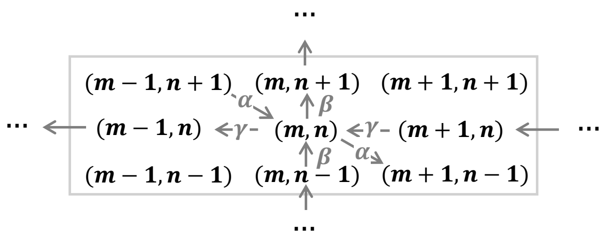


Fig. S3: State transitions in the trimolecular reaction.

The reactions can also be described by a continuous-time Markov chain, which is governed by the master equation S3. Assume the total number of the three molecules is $N$, and the numbers of molecule A and B are $m$ and $n$ respectively. Let $p(m, n, t)$ denote the probability of finding $m$ A and $n$ B simultaneously at time $t$.

The master equation for the system can then be written as:

$$\frac{\partial p(m,n,t)}{\partial t} = -p(m,n)[\hat{\alpha}mn + \hat{\beta}n(N-m-n) + \hat{\gamma}m(N-m-n)] +$$
$$\hat{\alpha}(m-1)(n+1)p(m-1,n+1) + \hat{\beta}(n-1)(N-m-n+1)p(m,n-1) +$$
$$\hat{\gamma}(m+1)(N-m-1-n)p(m+1,n) \quad (11)$$

where $(\hat{\alpha}, \hat{\beta}, \hat{\gamma}) = (\alpha, \beta, \gamma)/V$, $V$ is the volume of the reactor. We can then calculate the mean value of molecule number of A, B and C respectively. The dynamics of mean value of A can be obtained as:

$$\frac{\partial \langle m \rangle}{\partial t} = \sum_{m,n} m \frac{\partial p(m,n,t)}{\partial t} \quad (12)$$

Expanding the term $\partial p(m,n,t)/\partial t$ according to Eq. 25, we can write out the equation in a similar form as kinetic equation 8:

$$\frac{\partial \langle m \rangle}{\partial t} = \hat{\alpha}\langle mn \rangle - \hat{\gamma}\langle m(N-m-n) \rangle -$$
$$\left[ \sum_{m=N or n=0} \hat{\alpha}m(m+1)n + \sum_{n=N} \hat{\beta}mn(N-m-n) + \sum_{m=0} \hat{\gamma}m(m-1)(N-m-n) \right] p(m,n) \quad (13)$$

In above, The last term is a collection of third order moments, which is calculated based on the marginal distribution $(m, n = 0, N)$ and this term is exactly zero. Keeping only the first two terms and assuming that all elements in the covariance matrix between A and B concentration are negligible, we can get exactly the same equation as the kinetic equation. We can also write out the number dynamics of molecule B and C in a similar approach. If we keep the covariance terms in the equations, the set of ODEs does not generate sustained oscillation on ensemble average with the same parameters as in the kinetic equation. This suggests that there should be some other system coupled to the Markov chain to generate coherent oscillations.

Eq. 13 can be written in another form which includes explicitly individual terms of all moments:

$$\frac{\partial \langle m \rangle}{\partial t} = (\hat{\alpha} + \hat{\gamma})\langle mn \rangle - \hat{\gamma}N\langle m \rangle + \hat{\gamma}\langle m^2 \rangle \quad (14)$$

In order to close the equations we also need to write out dynamical equations for $\langle n \rangle, \langle m^2 \rangle, \langle n^2 \rangle, \langle mn \rangle, \ldots$. For approximation, we will simply cut off at second order moments and make additional assumptions about the third order moment as closure relations between third order moments and second order moments. Different versions can be obtained with different closure equations. The most general form up to second order moments can be written as:

$$\frac{d}{dt}\langle m\rangle = (\alpha+\gamma)\langle mn\rangle - \gamma N\langle m\rangle + \gamma\langle m^2\rangle \tag{15}$$

$$\frac{d}{dt}\langle n\rangle = -(\alpha+\beta)\langle mn\rangle + \beta N\langle n\rangle - \beta\langle n^2\rangle \tag{16}$$

$$\frac{d}{dt}\langle m^2\rangle = 2(\alpha+\gamma)\langle m^2 n\rangle + (\alpha-\gamma)\langle mn\rangle - \gamma(2N+1)\langle m^2\rangle + 2\gamma\langle m^3\rangle + \gamma N\langle m\rangle \tag{17}$$

$$\frac{d}{dt}\langle n^2\rangle = -2(\alpha+\beta)\langle mn^2\rangle + (\alpha-\beta)\langle mn\rangle + \beta(2N-1)\langle n^2\rangle - 2\beta\langle n^3\rangle + \beta N\langle n\rangle \tag{18}$$

$$\frac{d}{dt}\langle mn\rangle = (\alpha-\beta+\gamma)\langle mn^2\rangle + (-\alpha-\beta+\gamma)\langle m^2 n\rangle + (-\alpha+\beta N-\gamma N)\langle mn\rangle \tag{19}$$

In the main text, we use the "central-moment-neglect" approximation, also known as "zero-closure" [8]. An $m^{th}$ order "zero-closure" truncates the moment hierarchy by setting all central moments above order $m$ to zero. Specifically, in the second-order zero-closure scheme, all third-order central moments, such as $\langle(m-\langle m\rangle)^3\rangle$, are assumed to vanish.

### Supplementary note 5 - Derivation of governing principles for generating coherent oscillation in coupled Markov and non-Markov systems

#### Governing equations for coupled Markov and non-Markov system

In order to achieve coherent oscillation, we need to introduce a control input $u$, which is a function of the probability of each state. A general governing equation for the controlled system can be written as:

$$\frac{\partial \mathbf{P}}{\partial t} = \mathbf{M}_0\mathbf{P} + \mathbf{B}\mathbf{u} \tag{20}$$

There are numerous ways to design the control input $\mathbf{u}$. The most widely used methods include PID control, which refers to the proportional, integral, and derivative relationship between the control input $\mathbf{u}$ and system variable $\mathbf{P}$, respectively. It is clear that proportional control alone cannot generate oscillations. The reasoning is as follows: If We assume that $\mathbf{u} = \mathbf{K}\mathbf{P}$ and substitute it into Eq. 20, we obtain the modified governing equation:

$d\mathbf{P}/dt = (\mathbf{M}_0 + \mathbf{BK})\mathbf{P}$. In order to satisfy the probability conservation law ($\sum_{i=1}^{N} P_i = 1$), the matrix $\mathbf{M}_0 + \mathbf{BK}$ should also satisfy the condition that all rows sum to zero, as discussed in the previous section. However, this results in another Markov system, which does not generate coherent oscillations. Therefore, for the real biological system with coherent oscillations, the integral or derivative control mechanisms must be involved.

In this context, we propose a realization of integral control that generates coherent oscillations by coupling a stable Markov system to an unstable non-Markov system. This approach is inspired by multi-physics coupling observed in biological systems. For instance, the Hodgkin-Huxley model [9], which describes the initiation and propagation of action potentials in neurons, incorporates both ion channel dynamics-modeled as a Markov process-and membrane voltage dynamics, which are non-Markovian. Another example is the spontaneous oscillation of the basal surface contractions in a subset of follicle cells [10], where the behavior is governed by coupled mechano-chemical fields. The Hodgkin-Huxley model, in particular, serves as a prototypical example of a system combining Markovian and non-Markovian dynamics. The general form of the Hodgkin-Huxley model can be written as:

$$\frac{dV}{dt} = \left[-\bar{g}_{Na} m^3 h (V - V_{Na}) - \bar{g}_K n^4 (V - V_K) - g_L (V - V_L)\right]/C \quad (21)$$

$$\frac{dn}{dt} = \alpha_n(V)(1-n) - \beta_n(V)n \quad (22)$$

$$\frac{dm}{dt} = \alpha_m(V)(1-m) - \beta_m(V)m \quad (23)$$

$$\frac{dh}{dt} = \alpha_h(V)(1-h) - \beta_h(V)h \quad (24)$$

Here, $V$ is the membrane voltage, and $C$ is the membrane capacitance per unit area. $V_{Na}, V_K, V_L$ are the reversal potentials for sodium, potassium, and leak currents, respectively. $\bar{g}_i(i = Na^+, K^+, l)$ denotes the maximum conductance of sodium, potassium, and leak ion channels. The rate constants $\alpha_i, \beta_i$ describe the voltage-dependent transition rates of the $i$-th ion channel. The variables $n, m, h$ are dimensionless gating variables ranging between 0 and 1, representing the activation of potassium channel subunits, the activation of sodium channel subunits, and the inactivation of sodium channel subunits, respectively. The first equation describes the dynamics of the membrane voltage, treating the membrane as a capacitor and linking voltage changes to ion channel activity through voltage-sensitive opening probabilities. The remaining equations govern the dynamics of the gating variables, describing the fraction of open ion channels via a Markov process. These equations together form a canonical example of a coupled Markov and non-Markov system. If we discretize the gating variables $n, m, h$ and denote the joint state probability as $p(n, m, h, t)$, we can express the dynamics as a continuous-time Markov process with discrete states:

$$\begin{aligned}\frac{\partial p(n,m,h,t)}{\partial t} &= \alpha_n(V)(1-n+\Delta n)p(n-\Delta n,m,h,t)+\beta_n(V)(n+\Delta n)p(n+\Delta n,m,h,t)+\\ &\alpha_m(V)(1-m+\Delta m)p(n,m-\Delta m,h,t)+\beta_m(V)(m+\Delta m)p(n,m+\Delta m,h,t)+\\ &\alpha_h(V)(1-h+\Delta h)p(n,m,h-\Delta h,t)+\beta_h(V)(h+\Delta h)p(n,m,h+\Delta h,t)-\\ &[\beta_n(V)n+\alpha_n(V)(1-n)+\beta_m(V)m+\alpha_m(V)(1-m)+\beta_h(V)h+\alpha_h(V)(1-h)]p(n,m,h,t) \qquad (25)\end{aligned}$$

This equation, together with Eq. 21, forms a coupled Markov and non-Markov system capable of generating sustained and coherent voltage oscillations.

We can write out the most general form of the governing equation of the coupled system as:

$$\frac{\partial \mathbf{P}}{\partial t} = \mathbf{M_0}(\mathbf{x}) \cdot \mathbf{P} \qquad (26)$$

$$\frac{\partial \mathbf{x}}{\partial t} = f(\mathbf{P}, \mathbf{x}) \qquad (27)$$

In above, $\mathbf{P}$ is an $n$ dimensional vector that denotes the probability of each state in the Markov chain, while $\mathbf{x}$ is an $l$ dimensional vector which is the state variable of the coupled system (e.g., a mechanical system). These equations can be expanded up to the first order around a reference state-denoted as $\mathbf{P}_r$ and $\mathbf{x}_r$, typically corresponding to the steady state of the isolated Markovian and non-Markovian system -as follows:

$$\frac{\partial \mathbf{P}}{\partial t} = \left( \mathbf{M_0}(\mathbf{x}_r) + \left.\frac{\partial \mathbf{M_0}}{\partial \mathbf{x}}\right|_{\mathbf{x}=\mathbf{x}_r} \cdot \delta\mathbf{x} \right) \cdot (\mathbf{P}_r + \delta\mathbf{P}) \qquad (28)$$

$$\frac{\partial \mathbf{x}}{\partial t} = f(\mathbf{P}_r, \mathbf{x}_r) + \left.\frac{\partial f(\mathbf{P},\mathbf{x})}{\partial \mathbf{P}}\right|_{\mathbf{P}=\mathbf{P}_r,\mathbf{x}=\mathbf{x}_r} \cdot \delta\mathbf{P} + \left.\frac{\partial f(\mathbf{P},\mathbf{x})}{\partial \mathbf{x}}\right|_{\mathbf{P}=\mathbf{P}_r,\mathbf{x}=\mathbf{x}_r} \cdot \delta\mathbf{x} \qquad (29)$$

where $\mathbf{P} = \mathbf{P}_r + \delta\mathbf{P}, \mathbf{x} = \mathbf{x}_r + \delta\mathbf{x}$. In Eq. 28, $\frac{\partial \mathbf{M_0}}{\partial \mathbf{x}}$ is a third-order tensor, representing the derivative of matrix $\mathbf{M_0}$ with respect to vector $\mathbf{x}$. By neglecting second-order terms and rearranging the equations, we obtain the linearized governing equations as:

$$\frac{\partial \delta \mathbf{P}}{\partial t} = \mathbf{M_0}(\mathbf{x}_r) \cdot \delta \mathbf{P} + \left.\frac{\partial (\mathbf{M_0} \cdot \mathbf{P}_r)}{\partial \mathbf{x}}\right|_{\mathbf{x}=\mathbf{x}_r} \cdot \delta \mathbf{x} + \mathbf{M_0}(\mathbf{x}_r) \cdot \mathbf{P}_r \tag{30}$$

$$\frac{\partial \mathbf{x}}{\partial t} = \left.\frac{\partial f(\mathbf{P}, \mathbf{x})}{\partial \mathbf{P}}\right|_{\mathbf{P}=\mathbf{P}_r, \mathbf{x}=\mathbf{x}_r} \cdot \delta \mathbf{P} + \left.\frac{\partial f(\mathbf{P}, \mathbf{x})}{\partial \mathbf{x}}\right|_{\mathbf{P}=\mathbf{P}_r, \mathbf{x}=\mathbf{x}_r} \cdot \delta \mathbf{x} + f(\mathbf{P}_r, \mathbf{x}_r) \tag{31}$$

We further eliminate the constant terms through a variable substitution:

$$\tilde{\delta \mathbf{P}} = \alpha \delta \mathbf{P} + \mathbf{P}_{r2} \tag{32}$$

$$\tilde{\delta \mathbf{x}} = \alpha \delta \mathbf{x} + \mathbf{x}_{r2} \tag{33}$$

The shifting terms $\mathbf{P}_{r2}$ and $\mathbf{x}_{r2}$ are introduced to render equations 30-31 homogeneous, while the scaling factor $\alpha$ ensures that probabilities restored from the transformed variables ($\tilde{\delta \mathbf{P}}, \tilde{\delta \mathbf{x}}$) to the original variables ($\mathbf{P}, \mathbf{x}$ in Eqs.26–27) remain between 0 and 1. The resulting transformed homogeneous equations can be written as:

$$\frac{\partial \mathbf{P}}{\partial t} = \mathbf{M} \cdot \mathbf{P} + \mathbf{B}_1 \cdot \mathbf{x} \tag{34}$$

$$\frac{\partial \mathbf{x}}{\partial t} = \mathbf{B}_2 \cdot \mathbf{P} + \mathbf{C} \cdot \mathbf{x} \tag{35}$$

In the equations above, for simplicity, we replace $\tilde{\delta \mathbf{P}}, \tilde{\delta \mathbf{x}}$ by $\mathbf{P}$ and $\mathbf{x}$ respectively. Given the dimensions of $\mathbf{P} \in \mathbb{R}^n$ and $\mathbf{x} \in \mathbb{R}^l$, the coupling matrices $\mathbf{B}_1$ and $\mathbf{B}_2$ have dimensions $n \times l$ and $l \times n$. In order to satisfy probability conservation, $\mathbf{B_1}$ should satisfy: $\sum_i B_{1,ij} = 0$ for all $j$. Although the probabilities in the transformed homogeneous equations (Eqs. 34–35) still obey a conservation law, their sum is not necessarily equal to one. This discrepancy is resolved by reverting to the original equation.

In practice, when designing a coupled system to generate coherent oscillations, we begin with the transformed homogeneous system (Eqs. 34-35). The coefficient $\alpha$ in the variable substitution is introduced to ensure that the sum of probabilities across all states equals 1 in Eq. 28. The shifting terms $\mathbf{P}_{r2}$ and $\mathbf{x}_{r2}$ are introduced to ensure that the probabilities in Eq. 28 remain positive, maintaining the physical validity of the solution. An analytical solution of the coupled variable $x$ can be expressed in terms of probability $\mathbf{P}$ as:

$$\mathbf{x}(t) = e^{t\mathbf{C}}\mathbf{x}(0) + e^{t\mathbf{C}} \int_0^t e^{-s\mathbf{C}} \mathbf{B}_2 \mathbf{P}(s) ds \tag{36}$$

We can see that $\mathbf{x}$ is in the form of an integral of probability $\mathbf{P}$, which agrees with the design of an integral control.

#### Governing principles for generating coherent oscillation

We illustrate that with the coupling to another system, we can restore coherent oscillation. For this purpose, we seek to design the coupled system so that with the coupling matrices $\mathbf{B}_1, \mathbf{B}_2$, we are able to obtain at least one eigenvalue of the overall matrix with zero real part and non-zero imaginary part, which corresponds to coherent oscillation. To prevent instability, we also require that the real parts of other eigenvalues are non-positive. Denoting the new eigenvalues of the overall matrix ($\mathbf{A}$) as $\lambda$, we can obtain the characteristic equations as:

$$\det(\mathbf{A} - \lambda\mathbf{I}) = \det \begin{pmatrix} \mathbf{M} - \lambda\mathbf{I} & \mathbf{B}_1 \\ \mathbf{B}_2 & \mathbf{C} - \lambda\mathbf{I} \end{pmatrix} = 0 \tag{37}$$

Matrices $\mathbf{M}$ and $\mathbf{B}_1$ satisfy the condition that the sum of all rows equals zero. This is due to the conservation of probabilities. For convenience of calculating the determinant, we first perform some basic row and column matrix operations to remove this constraint.

$$\mathbf{A} - \lambda\mathbf{I} = \left(\begin{array}{cccc|ccc} M_{11}-\lambda & M_{12} & \dots & M_{1n} & B_{1,11} & \dots & B_{1,1l} \\ M_{21} & M_{22}-\lambda & \dots & M_{2n} & B_{1,21} & \dots & B_{1,2l} \\ \dots & \dots & \ddots & \dots & \dots & \ddots & \dots \\ M_{n1} & M_{n2} & \dots & M_{nn}-\lambda & B_{n,21} & \dots & B_{n,2l} \\ \hline B_{2,11} & B_{2,12} & \dots & B_{2,1n} & C_{11}-\lambda & \dots & C_{1l} \\ \dots & \dots & \ddots & \dots & \dots & \ddots & \dots \\ B_{2,l1} & B_{2,l2} & \dots & B_{2,ln} & C_{l1} & \dots & C_{ll}-\lambda \end{array}\right) \xrightarrow[r_i \to r_i + \frac{M_{i1}}{\lambda} r_1 (i=2,3,\dots,n)]{r_1 \to r_1 + \sum_{i=2}^{n} r_i} \tag{38}$$

$$\left(\begin{array}{cccc|ccc} -\lambda & -\lambda & \dots & -\lambda & 0 & \dots & 0 \\ 0 & M_{22}-M_{21}-\lambda & \dots & M_{2n}-M_{21} & B_{1,21} & \dots & B_{1,2l} \\ \dots & \dots & \ddots & \dots & \dots & \ddots & \dots \\ 0 & M_{n2}-M_{n1} & \dots & M_{nn}-M_{n1}-\lambda & B_{n,21} & \dots & B_{n,2l} \\ \hline 0 & B_{2,12}-B_{2,11} & \dots & B_{2,1n}-B_{2,11} & C_{11}-\lambda & \dots & C_{1l} \\ \dots & \dots & \ddots & \dots & \dots & \ddots & \dots \\ 0 & B_{2,l2}-B_{2,l1} & \dots & B_{2,ln}-B_{2,l1} & C_{l1} & \dots & C_{ll}-\lambda \end{array}\right) \xrightarrow{c_i \to c_i - c_1 (i=2,\dots,n)} \tag{39}$$

$$\left(\begin{array}{cccc|ccc} -\lambda & 0 & \dots & 0 & 0 & \dots & 0 \\ 0 & M_{22}-M_{21}-\lambda & \dots & M_{2n}-M_{21} & B_{1,21} & \dots & B_{1,2l} \\ \dots & \dots & \ddots & \dots & \dots & \ddots & \dots \\ 0 & M_{n2}-M_{n1} & \dots & M_{nn}-M_{n1}-\lambda & B_{n,21} & \dots & B_{n,2l} \\ \hline 0 & B_{2,12}-B_{2,11} & \dots & B_{2,1n}-B_{2,11} & C_{11}-\lambda & \dots & C_{1l} \\ \dots & \dots & \ddots & \dots & \dots & \ddots & \dots \\ 0 & B_{2,l2}-B_{2,l1} & \dots & B_{2,ln}-B_{2,l1} & C_{l1} & \dots & C_{ll}-\lambda \end{array}\right) \tag{40}$$

The new matrix has the same determinant as the original, which can be denoted as a block matrix $\mathbf{A}' - \lambda\mathbf{I}$:

$$\mathbf{A}' - \lambda\mathbf{I} = \begin{pmatrix} -\lambda & 0 & 0 \\ 0 & \mathbf{M}' - \lambda\mathbf{I} & \mathbf{B}'_1 \\ 0 & \mathbf{B}'_2 & \mathbf{C} - \lambda\mathbf{I} \end{pmatrix} = \begin{pmatrix} -\lambda & 0 \\ 0 & \mathbf{F} - \lambda\mathbf{I} \end{pmatrix} \tag{41}$$

We can clearly see that $\lambda = 0$ is an eigenvalue from the first element. Other eigenvalues can be obtained from the block matrix $\mathbf{F}$. In the new sub-matrix $\mathbf{F}$, there are no restrictions on the elements in matrix $\mathbf{B}'_1$. Assume that $\mathbf{C} - \lambda\mathbf{I}$ is invertible, the characteristic equation corresponding to matrix $\mathbf{F}$ can then be written as: $\det(\mathbf{F} - \lambda\mathbf{I}) = \det(\mathbf{C} - \lambda\mathbf{I}) \det[(\mathbf{M}' - \lambda\mathbf{I}) - \mathbf{B}'_1(\mathbf{C} - \lambda\mathbf{I})^{-1}\mathbf{B}'_2] = 0$. The matrix $\mathbf{M}'$ can be transformed to a quasi upper triangular matrix via Schur decomposition: $\mathbf{M}' = \mathbf{P}_1\mathbf{D}_1\mathbf{P}_1^{-1}$. Here $\mathbf{P}_1$ is a unitary matrix which satisfies that $\mathbf{P}_1\mathbf{P}_1^T = \mathbf{I}$. $\mathbf{D}_1$ is a quasi upper triangular matrix with square blocks along the diagonal, where all entries below the subdiagonal are zero. We denote the diagonal blocks as $D_{1,ii}$, which are either scalars($d_{1,ii}$) or 2×2 matrices. The eigenvalues of all the diagonal blocks are the eigenvalues of matrix $\mathbf{M}'$. Here we assume that there are totally $n_b$ blocks, in which there are $p$ single-element blocks ($p \leq n_b$). All single element blocks are positioned at the top left of the matrix $\mathbf{D}_1$ with a descending order $(d_{1,11} > d_{1,22} > \dots)$. The indices satisfy the identity: $p + 2(n_b - p) = n - 1$. These single elements are exactly the eigenvalues of matrix $\mathbf{D_1}$, which can be denoted as $\sigma_i^{(1)}$. According to our definition, all $\sigma_i^{(1)}$ with $i \leq p$ are real, and they are complex when $i > p$. The rest of the diagonal elements are all 2 by 2 matrices, which can be denoted in the capital letter $D_{1,ii}, i = p+1, \dots, n_b$. Assume that the matrix of the coupled system is diagonalizable. Matrix $\mathbf{C}$ can then be decomposed as: $\mathbf{C} = \mathbf{P}_2\mathbf{D}_2\mathbf{P}_2^{-1}$, where $\mathbf{D}_2 = \text{diag}(\sigma_1^{(2)}, \sigma_2^{(2)}, ..., \sigma_l^{(2)})$ are diagonal matrices of the eigenvalues of the coupled system $\mathbf{C}$, and $\mathbf{P}_2$ is the eigenvector matrix whose columns are eigenvectors corresponding to the eigenvalue matrix. After diagonalization, the characteristic equation corresponding to matrix $\mathbf{F}$ can be rewritten as:

$$\det[\mathbf{P}_1(\mathbf{D}_1 - \lambda\mathbf{I})\mathbf{P}_1^{-1} - \mathbf{B}_1'\mathbf{P}_2(\mathbf{D}_2 - \lambda\mathbf{I})^{-1}\mathbf{P}_2^{-1}\mathbf{B}_2'] = 0 \tag{42}$$

$$\det[(\mathbf{D}_1 - \lambda\mathbf{I}) - \mathbf{P}_1^{-1}\mathbf{B}_1'\mathbf{P}_2(\mathbf{D}_2 - \lambda\mathbf{I})^{-1}\mathbf{P}_2^{-1}\mathbf{B}_2'\mathbf{P}_1] = 0 \tag{43}$$

Let $\mathbf{Q}_1 = \mathbf{P}_1^{-1}\mathbf{B}_1'\mathbf{P}_2$ and $\mathbf{Q}_2 = \mathbf{P}_2^{-1}\mathbf{B}_2'\mathbf{P}_1$, the characteristic equation can then be written as:

$$\det[(\mathbf{D}_1 - \lambda\mathbf{I}) - \mathbf{Q}_1(\mathbf{D}_2 - \lambda\mathbf{I})^{-1}\mathbf{Q}_2] = 0 \tag{44}$$

It is impractical to obtain general analytical expressions for the eigenvalues of the new matrix $\mathbf{F}$. However, with certain restrictions on matrix $\mathbf{Q}_1$ and $\mathbf{Q}_2$, we are able to obtain analytical results. A useful simplification is to assume that matrix $\mathbf{Q}_1$ and $\mathbf{Q}_2$ are diagonal. The underlying assumption is that each degree of freedom in the non-Markov system corresponds to each degree of freedom in the Markov system in a one-to-one mapping, without any additional crosstalk. Note that in general, $\mathbf{Q}_1$ and $\mathbf{Q}_2$ are rectangular, diagonal here means all the entries not of the form $Q_{ii}$ being zero. The diagonal elements of the two matrices are denoted as: $q_i^{(1)}, q_i^{(2)} (i = 1, 2, ..., min\{n-1, l\})$. With the diagonal assumption, we can further write the characteristic equation as:

$$\prod_{i \in S_1} \left( \sigma_i^{(1)} - \lambda - \frac{q_i^{(1)} q_i^{(2)}}{\sigma_i^{(2)} - \lambda} \right) \prod_{j \in S_2} \det \left[ D_{1,jj} - \lambda\mathbf{I} - \text{diag} \left( \frac{q_\alpha^{(1)} q_\alpha^{(2)}}{\sigma_\alpha^{(2)} - \lambda}, \frac{q_\beta^{(1)} q_\beta^{(2)}}{\sigma_\beta^{(2)} - \lambda} \right) \right] = 0 \tag{45}$$

Here $S_1$ is the set of indices corresponding to scalar diagonal elements in matrix $\mathbf{D}_1$, which is simply $S_1 = 1, 2, \dots, p$. $S_2$ is the set of indices corresponding to $2{\times}2$ blocks in $\mathbf{D}_1$, which is $S_2 = \{p+1, \dots, n_b\}$. $\alpha, \beta$

are the indices of individual scalar entries corresponding to the $j^{th}$ block. In our definition of the matrix, $\alpha = 2j - p - 1, \beta = 2j - p$. When the dimension of the matrix of non-Markov system is smaller than that of the Markov system ($l < n - 1$), there is no definition of $q_\alpha, q_\beta$ when $\alpha, \beta > n - 1$. In such a case, we assume that $q_\alpha = q_\beta = 0$. Notice that under our theoretical framework, the design is meaningful only when $l \leq n - 1$. When the dimension of the non-Markov system is higher than that of the Markov system, the eigenvalues of the non-Markov system matrix with indices larger than $n - 1$ will not contribute to the coupled whole system.

From Eq. 45, we identify two distinct approaches for achieving purely imaginary eigenvalues, which correspond to coherent oscillations. The first approach targets the real eigenvalues of the Markov system (first term in Eq. 45), while the second targets complex eigenvalues (second term; see Fig. **??**). These strategies induce different dynamical transitions: the first converts a stable, non-oscillatory mode into a coherent oscillation, whereas the second stabilizes an otherwise decaying oscillatory mode. When targeting the real eigenvalues (first approach), we obtain the following equations:

$$\lambda^2 - (\sigma_i^{(1)} + \sigma_i^{(2)})\lambda + (\sigma_i^{(1)}\sigma_i^{(2)} - q_i^{(1)}q_i^{(2)}) = 0, i \in S_1 \tag{46}$$

Here we call $\sigma_i^{(1)}$ target eigenvalue of the Markov system, which corresponds to a specific mode of the solution top the original Markov system. Here our target eigenvalues $\sigma_i^{(1)}$ are all real, and it corresponds to a decaying mode without oscillation. Otherwise, it corresponds to a decaying mode with oscillation. We can see that Eq. 46 is a quadratic function with real coefficients. In order to generate coherent oscillation, we require that there exist eigenvalues with zero real part, which gives the condition that one of the eigenvalues of the coupling system $\sigma_i^{(2)}$ should satisfy $\sigma_i^{(1)} + \sigma_i^{(2)} = 0$ and $\sigma_i^{(1)}\sigma_i^{(2)} - q_i^{(1)}q_i^{(2)} > 0$. For other eigenvalues, the real part of $\lambda$ should be non-positive. The angular frequency of the coupled system is $\omega = \sqrt{\sigma_i^{(1)}\sigma_i^{(2)} - q_i^{(1)}q_i^{(2)}}$.

The condition for coherent oscillation can be interpreted as follows: Since the Markov system always have eigenvalues with negative real parts, $\sigma_i^{(1)} < 0$ always holds. In order to satisfy the condition above, $\sigma_i^{(2)} > 0$ must hold, which means the coupling system should be inherently unstable. Another requirement that $\sigma_i^{(1)}\sigma_i^{(2)} - q_i^{(1)}q_i^{(2)} > 0$ means that $-q_i^{(1)}q_i^{(2)}$ should be large enough to make the whole term positive. In order to achieve this, we require that: 1. There should be a strong coupling between the two systems. 2. The dependence of one system on the other should be opposite the other way round (e.g., System A inhibits B while B enhances A). This requirement resembles a negative feedback system.

When the non-Markov system is targeting the complex eigenvalues of the Markov matrix, the second product in Eq. 45 should generate pure imaginary roots. We denote individual elements in matrix $D_{1,ii}$ as $d_{\alpha\beta}$ ($i \in S_2$). The new eigenvalues $\lambda$ of the coupled system can then be solved from the following equation:

$$\left[d_{\alpha\alpha}^{(1)} - \lambda - \frac{q_\alpha^{(1)} q_\alpha^{(2)}}{\sigma_\alpha^{(2)} - \lambda}\right]\left[d_{\beta\beta}^{(1)} - \lambda - \frac{q_\beta^{(1)} q_\beta^{(2)}}{\sigma_\beta^{(2)} - \lambda}\right] - d_{\alpha\beta}^{(1)} d_{\beta\alpha}^{(1)} = 0 \tag{47}$$

The equation above is a quartic equation which generates four roots. For simplicity, we further assume that the $\beta^{th}$ element in the non-Markov system is weakly coupled to the Markov system. We can then make the approximation that $q_\beta^{(1)} = 0$. This approximation will decouple one dimension from the quartic equation, thus simplifying the quartic equation to a cubic equation:

$$\lambda^3 - \left(d_{\alpha\alpha}^{(1)} + d_{\beta\beta}^{(1)} + \sigma_\alpha^{(2)}\right)\lambda^2 + \left[\sigma_\alpha^{(2)}\left(d_{\alpha\alpha}^{(1)} + d_{\beta\beta}^{(1)}\right) + d_{\alpha\alpha}^{(1)} d_{\beta\beta}^{(1)} - d_{\alpha\beta}^{(1)} d_{\beta\alpha}^{(1)} - q_\alpha^{(1)} q_\alpha^{(2)}\right]\lambda + \left[q_\alpha^{(1)} q_\alpha^{(2)} d_{\beta\beta}^{(1)} - \sigma_\alpha^{(2)}\left(d_{\alpha\alpha}^{(1)} d_{\beta\beta}^{(1)} - d_{\alpha\beta}^{(1)} d_{\beta\alpha}^{(1)}\right)\right] = 0 \tag{48}$$

The equation above is in the form $a\lambda^3 + b\lambda^2 + c\lambda + d = 0$. To ensure that the equation contains pure imaginary root, the equation can be factorized as: $(\lambda + r)(\lambda^2 + \alpha^2) = 0$. We can see that a necessary and sufficient condition for existence of pure imaginary root is that $bc = ad$ and $b > 0, c > 0$. Other criterion can also be explored. For example, to ensure that the equation contains oscillatory mode (the equation contains complex roots), the discriminant needs to satisfy: $\Delta = b^2c^2 - 4ac^3 - 4b^3d - 27a^2d^2 + 18abcd < 0$. To ensure stability, the necessary and sufficient condition is: $a, b, c, d > 0, \quad bc - ad > 0$, according to the Hurwitz criterion.

Notably, in this work, "sustained oscillation" refers to the ensemble-average behavior, characterized by both persistent amplitude and temporal coherence in a single trajectory, where coherence is quantified by the decay rate of the autocorrelation function's amplitude. In the main text, we demonstrated that the average time dynamics exhibit sustained oscillations when the system is coupled to a non-Markovian component. Here, we additionally compute the autocorrelation function for both the isolated and coupled systems in terms of the number of molecules of species A ($n_A$). The auto-correlation function can be obtained as the following:

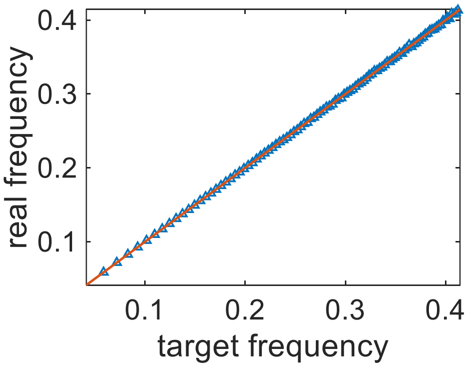


Fig. S4: Comparison between target frequency and real frequency by designing.

$$C_A(t) = \langle n_A(0) n_A(t) \rangle \tag{49}$$

Where the average is taken over different realizations (ensemble average). This quantity can be evaluated from the initial probability density ($P(\mathbf{n}, 0)$) and the conditional probability density at time $t$ ($P(\mathbf{n}', t|\mathbf{n}, 0)$) given the initial molecule number $\mathbf{n} = (n_A, n_B, n_C)$ as:

$$C_A(t) = \sum_{\mathbf{n}'} \sum_{\mathbf{n}} n_A(0) n_A(t) P(\mathbf{n}, 0) P(\mathbf{n}', t|\mathbf{n}, 0) \tag{50}$$

The autocorrelation function decays at the same rate as the probability density function, as shown in [11]. Therefore, the non-decaying property of the ensemble-averaged dynamics directly implies non-decaying temporal coherence. This is illustrated in Fig. 3c, where we compare the autocorrelation functions of the isolated and coupled systems. Coupling to a non-Markovian component preserves both oscillation amplitude and temporal coherence, as evidenced by the non-decaying autocorrelation function.

Our model provides a theoretical framework for designing sustained coherent oscillators with a desired frequency and various oscillation modes. Fig. S4 demonstrates the accuracy of our design within a target frequency. When targeting the oscillatory decaying mode, similar to the directly decaying mode, we obtain a phase diagram showing a square-root (1/2) scaling of the oscillation frequency with coupling strength between the Markovian and non-Markovian systems near the coherent oscillation regime (Fig. S5). Multiple frequency components can be generated by coupling to different modes in the Markov system, giving various wave shapes. However, as the system dimension increases, the predictions become more sensitive to noise and perturbations. The larger matrices involved lead to higher errors in eigenvalue calculations, impacting accuracy of results (Fig. S6).

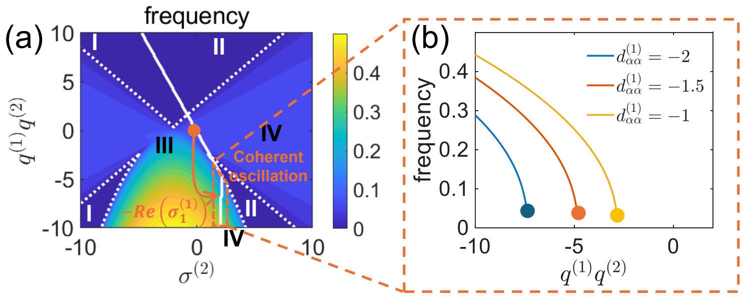


Fig. S5: (a) Phase diagram of the oscillation frequency as a function of the non-Markov eigenvalue ($\sigma^{(2)}$) and coupling strength ($q^{(1)}q^{(2)}$), targeting an oscillatory Markov mode. (b) Phase transition of the oscillation frequency versus coupling strength when targeting an oscillatory decaying Markov mode, exhibiting square-root (1/2) scaling behavior.

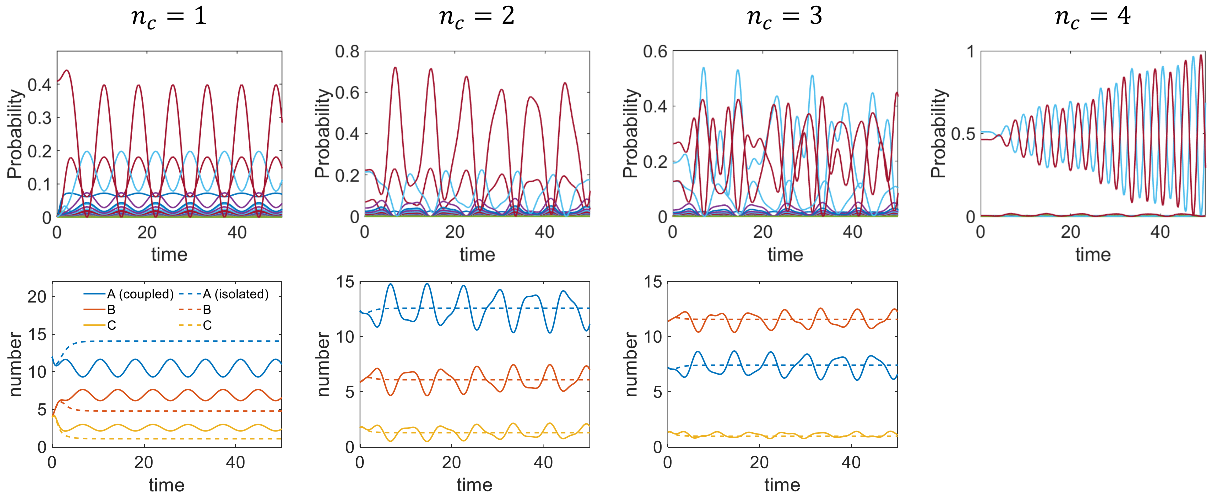


Fig. S6: Time evolution of the state probabilities with varying dimensions of the non-Markov system. Combining different frequency components generates more complex waveforms. However, increasing the dimension of the non-Markov system may lead to instability. $n_c$ represents the number of target eigenvalues (real eigenvalues), which corresponds to the coupled degrees of freedom between the Markov system and the non-Markov system.
